# PCAGO: An interactive tool to analyze RNA-Seq data with principal component analysis

**DOI:** 10.1101/433078

**Authors:** Ruman Gerst, Martin Hölzer

**Affiliations:** Applied Systems Biology, Leibniz Institute for Natural Product Research and Infection Biology, HKI, Jena, Germany; Faculty of Biological Sciences, Friedrich Schiller University Jena, Jena, Germany; RNA Bioinformatics and High-Throughput Analysis, Friedrich Schiller University Jena, Leutragraben 1, Jena, Germany; European Virus Bioinformatics Center, Friedrich Schiller University Jena, Leutragraben 1, Jena, Germany

## Abstract

The initial characterization and clustering of biological samples is a critical step in the analysis of any transcriptomics study. In many studies, principal component analysis (PCA) is the clustering algorithm of choice to predict the relationship of samples or cells based solely on differential gene expression. In addition to the pure quality evaluation of the data, a PCA can also provide initial insights into the biological background of an experiment and help researchers to interpret the data and design the subsequent computational steps accordingly. However, to avoid misleading clusterings and interpretations, an appropriate selection of the underlying gene sets to build the PCA and the choice of the most fitting principal components for the visualization are crucial parts. Here, we present PCAGO, an easy-to-use and interactive tool to analyze gene quantification data derived from RNA sequencing experiments with PCA. The tool includes features such as read-count normalization, filtering of read counts by gene annotation, and various visualization options. In addition, PCAGO helps to select appropriate parameters such as the number of genes and principal components to create meaningful visualizations.

**Availability and implementation:** PCAGO is implemented in R and freely available at github.com/hoelzer-lab/pcago. The tool can be executed as a web service or locally using a Docker image.

## Introduction

In the last decade, the sequencing of whole transcriptomes (RNA-Seq) emerged as a powerful technique to understand versatile molecular mechanisms and to tackle various biological questions. The initial characterization and clustering of biological samples based on their different gene expression levels are crucial steps in the early access of RNA-Seq experiments. Principal component analysis (PCA)^1^ is a technique used to emphasize variation and to carve out strong patterns in a high-dimensional data set, such as a matrix of gene expression val ues^2^. The output of a PCA transformation can be used to make data easy to explore and visualize. PCAs are therefore widely used approaches in bioinformatics^3^ and should be an important part of almost all RNA-Seq analysis pipelines today^4^.

Although PCAs are simple to compute and frequently used in transcriptomics, metabolomics, and medical studies, there are several challenges and disadvantages that can especially occur when analyzing experimental data^5^. For example, the visualization and interpretation of the data can be heavily influenced by the considered principal components, the number of most-variant genes and prior normalization of gene expression^2,5^. Typically, the main components that contribute most to the total variance deviation are selected. In many cases, however, interesting results cannot be observed directly by focusing only on the components with the greatest variance, e.g., if the variances of the individual components are close to each other. Therefore, a careful evaluation of the PCA and its components is crucial to derive meaningfully interpretations from the transformation.

In general, RNA-Seq data are of high dimensionality as they are based on the expression of thousands of genes in a much smaller set of samples^3^. A low-dimensional projection can be achieved with singular value decomposition such as a PCA. The data is presented in the form of a smaller number of variables called principal components (PCs) constructed by performing singular value decompositions of the original variables (one read count per gene and sample). In other words, a PCA transforms quantified expression information from correlated genes (represented by a matrix of read counts) into PCs or, when used in gene expression analysis, into ‘metagenes’^6^. PCs or ‘metagenes’ can be considered as a group of genes with correlated expression. The PCs are orthogonal to each other and can be plotted to represent the grouping of the measured expression data. For a more detailed description and formal mathematical definition of a PCA in the context of gene expression analysis, see^3^. As a best practice, a PCA should first be used for quality control, e.g., to check whether biological replicates have a similar expression profile and tend to cluster together^4^.

By definition, the first principal component (PC1) explains the greatest possible variance in the data (e.g., for an infection experiment, 40% of the variance between control and infected samples is explained by the genes that define PC1). PC2 explains the second greatest portion of variance, and so on. Traditionally, the first few PCs are used for visualization since they capture most of the variation from the original data set.

However, less informative PCs such as PC3 or PC4 might capture strong patterns comparable to PC1 and PC2 and help to distinguish between gene expression patterns more clearly^2^. Different gene sets that can be used as the backbone of the PCA, as well as a varying number of PCs, can influence the effectiveness in capturing the cluster structure^2,3^. Thus, low-variance PCs might be also usable to identify ubiquitously expressed genes (e.g., potential house-keeping genes), which could then be used as an additional filtering step to define a more reliable set of genes as the backbone of PCA visualization. Accordingly, there is a great need to investigate the effectiveness of the PCA transformation before drawing conclusions on the gene expression data^2^.

Here, we present PCAGO, an interactive and easy-to-use tool to analyze and visualize quantification data derived from RNA-Seq with PCA. A web service allows the direct execution of PCAGO, or, especially for local use, the tool can be started with a single command from a Docker container. The functionality can be easily tested by loading sample RNA-Seq read counts obtained from Klassert et al.^7^ and Riege et al.^8^ (execute *Start analysis* → *Load all sample data*).

In contrast to other PCA tools^9,10^, PCAGO gives the user the opportunity to comprehensively inspect and vary key features of the transformation, such as the PCs, and to easily compare different outcomes. Furthermore, PCAs can be built on annotation-filtered gene subsets, that might better reflect the biological question of interest in the final visualization. A minimum set of genes that produces the same clustering as all genes can be calculated and an animated PCA shows how PCs and the clustering change by incrementally including less variant genes in the transformation. Finally, all computational steps are well documented to allow full reproducibility of the performed PCA.

## PCAGO workflow and features

PCAGO requires a table of raw or already normalized read count data as produced by any standard RNA-Seq pipeline^4^ as input (Fig. 1A). Based on the raw read counts, PCAGO can perform the following steps: normalization (DESeq2-based^11^, TPM^12^); sample and gene set annotation; Ensembl and gene ontology (GO) integration from online databases; filtering of gene subsets by gene annotation and variance cutoff; visualization of dendrograms, 2D/3D-PCAs, scree plots, and PC loadings; PCA plot animation; export of all results in various formats; and the creation of an HTML-based documentation for full reproducibility. While most of these steps are optional, at least one read count table and corresponding sample annotation must be provided to perform the PCA (Fig. 1A). All output of PCAGO can be downloaded in various formats (pdf, svg, png, tiff, mp4) and is directly visualized in the application (Fig. 1B).

**Figure 1:**
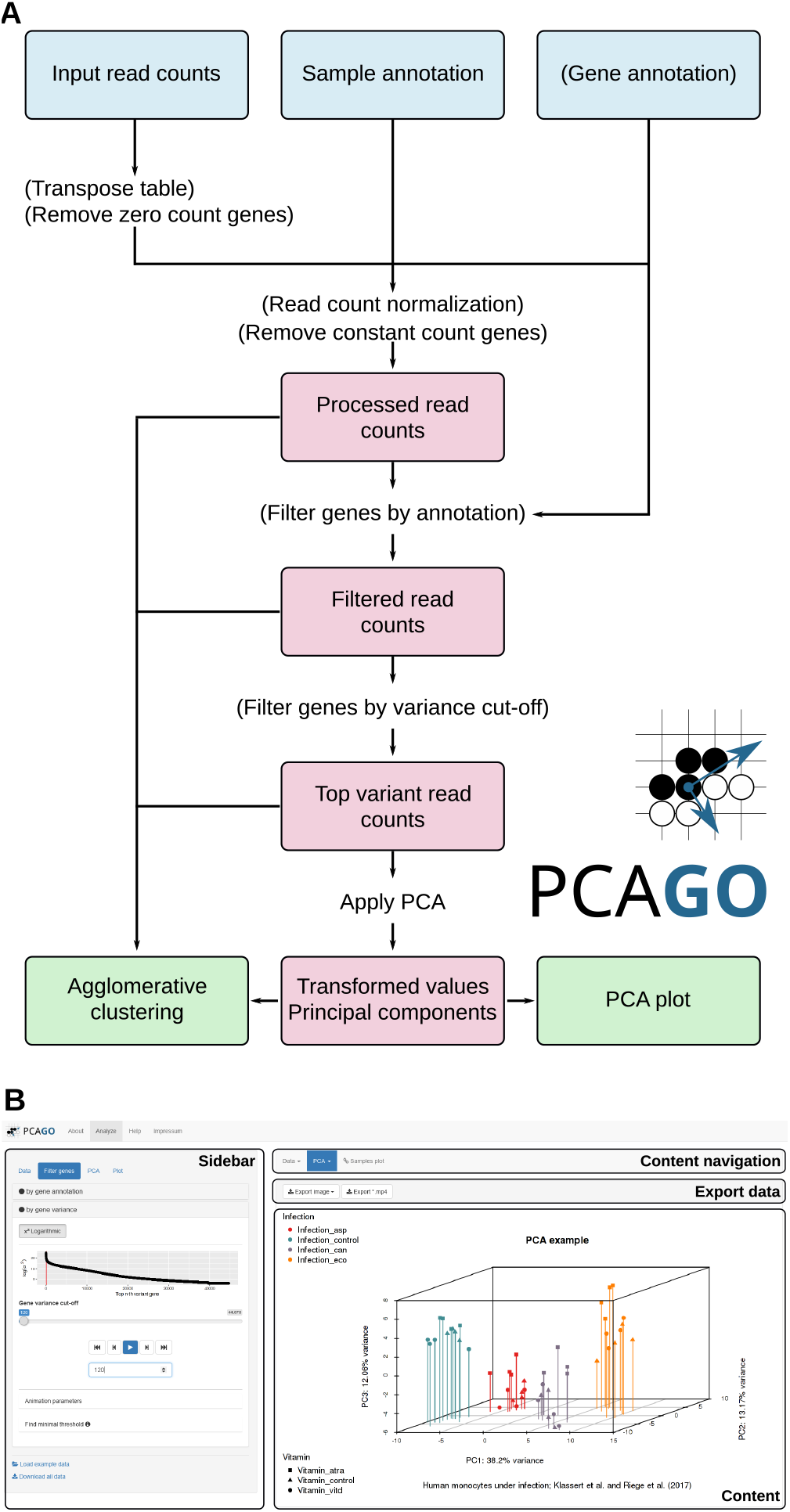
**(A)** Workflow of PCAGO. Optional steps are written in brackets. **(B)** Interface of the application. Example data^7,8^ was loaded and a 3D-PCA plot drawn.

Based on microarray-derived expression data, De Haan et al.^13^ showed the advantages of performing PCAs only on certain gene subsets. One of the major used methods to categorize functional gene information is the Gene Ontology (GO) database^14^. With PCAGO, different gene subsets filtered by features such as chromosome, Ensembl^15^ biotype (protein-coding, miRNA, lncRNA,…) or GO terms can be defined and directly used for the PC calculations. An annotation can be applied by the user or directly extracted from online databases such as Ensembl BioMart^16^.

The heart of PCAGO is the visualization of interactive 2D/3D-PCA and hierarchical clustering plots as well as scree and loading plots of the PCs (see online manual). An animation feature is included to analyze how adding more and more variant genes into the calculation changes the outcome. We included highly flexible customization and coloring options directly within the application to generate publication-ready vector graphics. All generated data, as well as detailed reports about the processing steps, can be viewed and exported in various formats. As a special feature, PCAGO can export an MP4 movie of the animated PCA plot.

Only recently, Boileau *et al.* presented scPCA, sparse contrastive PCA, to explore highdimensional biological data^17^. They show that scPCA can produce denser and more relevant clusters than for example t-SNE or UMAP, thus, resulting in more easily interpretable PC structures. We plan to integrate their methodology in a next release of PCAGO.

## Availability and implementation

PCAGO is implemented in R18 and distributed on the powerful R Shiny framework. The application, including a comprehensive online documentation with how-to and help sections, as well as an easy-to-load example data set, is freely available at pcago.bioinf.unijena.de. However, we strongly recommend using a Docker container that can be easily started with a single command: `docker run -p 8000:8000 --rm-it mhoelzer/pcago:latest./run_packrat.sh` Besides, an independent desktop version developed with Electron is also available in the GitHub repository. The source code, as well as a list of all used software packages, are available on GitHub (github.com/hoelzer-lab/pcago). PCAGO can be also installed and run locally, based on a working R (v3.4.3 or higher) and MariaDB installation.

## Acknowledgement and funding

MH has been supported by the Deutsche Forschungsgemeinschaft (DFG) through the Priority Program SPP-1596 MA5082/7-1 and DFG CRC 1076 “AquaDiva”, subproject A06. RG is funded by the International Leibniz Research School for Microbial and Biomolecular Interactions (ILRS). Special thanks go to Lasse Faber for the realisation of the Docker image.

## Author Contributions

MH developed the research idea. RG did the methods development and implementation of the tool. RG implemented all functionalities of the web service. MH wrote the main manuscript. MH and RG proofread and finalized the manuscript for publication.

